# The Rationalization of Carbon Monoxide and Hemoglobin Association

**DOI:** 10.1101/2025.05.27.656340

**Authors:** Chihjen Lee, Nikki Chen

## Abstract

Carbon monoxide is a colorless, odorless, and poisonous gas, responsible for approximately 100,000 emergency room visits and over 420 deaths in the U.S. each year. Both carbon monoxide and oxygen bind to the ferrous ions in hemoglobin, but carbon monoxide has a significantly higher affinity. Extensive research has been conducted on the interaction between carbon monoxide and hemoglobin. However, a straightforward and practically applicable equation describing the relationship between carbon monoxide saturation and pressure is not found in the existing literature.

In this paper, we establish an equation and confirm that the plot of CO saturation against CO pressure follows a hyperbolic shape, characterized by a continuous decrease in slope. In contrast, the oxygen-hemoglobin association curve is sigmoidal. These distinct curve shapes have different physiological implications. Our equation enables the determination of one variable—either saturation or pressure—if the other is known.

Further analysis reveals the distribution of all five species of carboxyhemoglobin, showing that the triply bound form is abundant—a notable contrast to the distribution of oxyhemoglobin species. Additionally, our equation confirms that carbon monoxide’s affinity for hemoglobin is approximately 230 times higher than that of oxygen. Lastly, we propose a new general equation that may generate all carbon monoxide-hemoglobin association curves under various oxygen pressures.

**New and Noteworthy:** In this paper, we introduce the first known set of equations that model the Carbon Monoxide Hemoglobin association curve and the Carbon Monoxide Hemoglobin Fractional association curves. These novel equations represent a noteworthy advancement in the ability to quantitatively model CO binding to hemoglobin. We present the methodology utilized that explains the derivation of precise coefficients for these equations, along with the data points used to calculate the coefficients. Unlike existing models, our equations allow the direct computation of the CO saturation from the CO association equation and the CO fraction association equations from the partial pressure of carbon monoxide. This approach simplifies the mathematical modeling process while preserving a high degree of biological significance. Furthermore, the equations are built upon a complex and precise methodological foundation, enabling a deeper understanding of the binding behavior of hemoglobin in response to varying CO levels. This quantitative model has the potential to be applied to clinical practices in medicine. It provides insight into the complex relationship between CO and hemoglobin that is critical to many biological processes.

## Introduction

A form of the Coburn–Forster–Kane equation (CFKE) is expressed as a differential equation that relates the rate of change of carbon monoxide (CO) saturation to the mean capillary CO pressure.

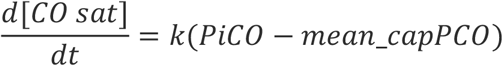

The solution to this equation would express CO saturation as a function of time. In contrast, the goal of our paper is to express CO saturation as a function of CO pressure. In clinical practice, both arterial CO saturation and CO pressure can be measured in a laboratory. Alternatively, a pulse CO-oximetry can be used to monitor CO saturation continuously without a blood test. With the help of our equation, arterial PCO can then be quickly determined.

In 2023, Chou and Lee developed a theory of oxygen-hemoglobin association and derived an equation based on clinical data sets [Chou and Lee, 2023]. Their association equation was found to be a rational function of the fourth degree. In this paper, we aim to establish the carbon monoxide hemoglobin association equation in a similar manner. We demonstrate that the CO association equation is also a rational function of the fourth degree, but it exhibits a different shape when plotted. At an oxygen pressure of 100 mmHg, a simplified version of the equation is presented below, PCO in ppt:

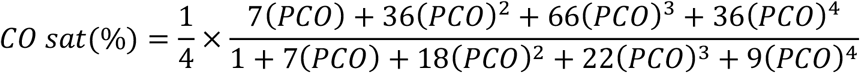

Both carbon monoxide and oxygen bind to the ferrous ion of the hemoglobin molecule. In normal atmospheric conditions, the concentration of carbon monoxide is minimal (approximately 100 ppb), resulting in a hemoglobin saturation of only 0.02% [Hatheway et al., 2023]. Consequently, carbon monoxide was considered negligible in the oxygen-hemoglobin association equation. However, deriving the CO hemoglobin association equation requires accounting for both carbon monoxide and oxygen, adding complexity to the equation’s formulation.

Assuming an equilibrium oxygen pressure of 100 mmHg, we first establish the CO saturation equation by considering all four oxyhemoglobin and carboxyhemoglobin species, their various combinations, and pure hemoglobin molecules. Next, using data from literature, we obtain four equilibrium data points for CO pressure and CO saturation. This allows us to solve for four unknowns using four independent linear equations [Chou and Lee, 2023]. The equation is then simplified by rounding off the coefficients. A step-by -step derivation is presented in the following section.

## Methods

### Deriving the Carbon monoxide hemoglobin association equation

Each hemoglobin molecule contains four heme groups and, consequently, four ferrous ions. Each ferrous ion can bind to either O_2_ or CO, allowing each hemoglobin molecule to bind up to four molecules simultaneously. As a result, the association of hemoglobin with oxygen and carbon monoxide can produce the following species:

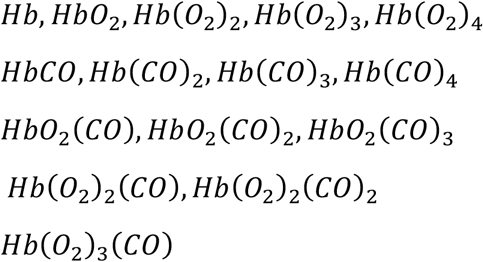

A general notation reaction is shown below, where *i* represents the number of O_2_ molecules bound and *j* represents the number of CO molecules bound. Notably, *i, j*, and *i* + *j* all range from 0 to 4, as each hemoglobin molecule has four binding sites.

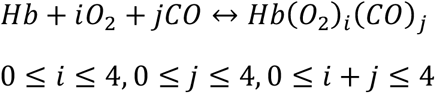

According to laws of chemical kinetics, we can derive equation (1) as shown below.

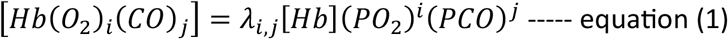

*λ*_*i,j*_ is the association constant.

Carbon monoxide saturation is defined as the ratio of CO molecules bound to hemoglobin to the total possible CO binding sites, as represented by the equation below.

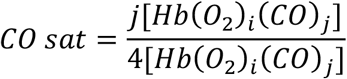

By substituting [*Hb*(*O*_2_)_*i*_(*CO*)_*j*_] with λ_*i,j*_[*Hb*](*PO*_2_)^*i*^(*PCO*)^*j*^ as shown in equation (1), we get equation (2):

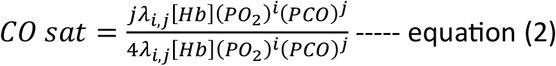

By canceling out [Hb], we get:

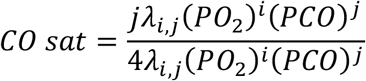

We rewrite the above equation by replacing *j* with all its potential values (0 ≤ *j* ≤ 4): *CO sat*

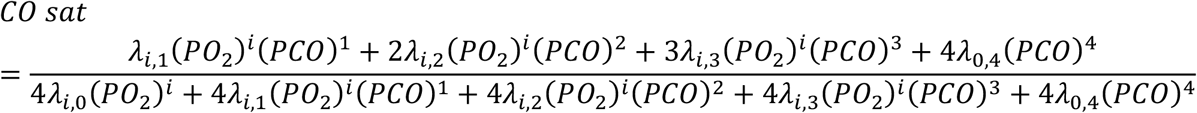

We then factor out and divide the numerator and denominator by λ_*i*,0_(*PO*_2_)^*i*^.

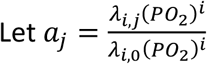

With substitution, the equation can be rewritten as:

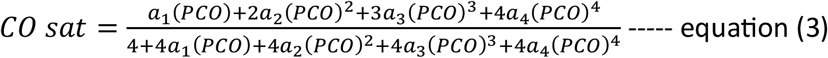

Please note that in this case, PO_2_ equals 100, and λ_*i,j*_ is constant. Therefore, *a*_*j*_ represents the coefficients when the oxygen pressure is 100 mmHg. However, *a*_*j*_ would have different values at varying PO_2_ levels or oxygen pressures, thereby modifying the CO saturation equations and causing a “rightward” or “leftward” shift of the curves.

Listed below are some of the coefficients:

1. λ_0,0_ = 1
2. *a*_0_ = 1
3. *a*_*j*_ is a function of *PO*_2_. Therefore, the carbon monoxide hemoglobin association curve “shifts” to right or left, depending on *PO*_2_ values
4. 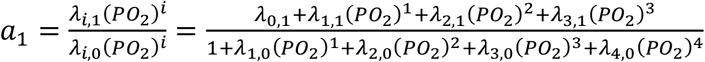
5. 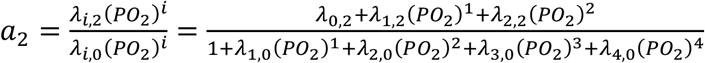
6. 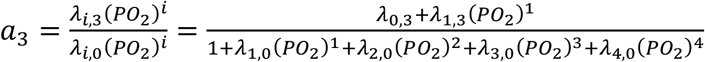
7. 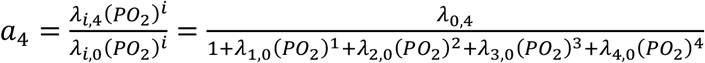

Equation (3) is solvable when four (PCO, CO saturation) data points are provided, as established by Chou and Lee [Chou and Lee, 2023]. From the literature, we identified the following equilibrium data points for (PCO, CO saturation) as reported by Hess et al. [Hess et al., 2017]. The pressure unit for CO is in ppt (parts per thousand).

(0.0087, 1.71%), **(0.025, 4.21%)**, (0.035, 5.82%), (0.050, 7.79%), **(0.1, 14.83%)**, (0.2, 25.96), **(0.5, 46.61%), (1, 63.58%)**

We can substitute those four bolded data points into Equation (3) and solve for *a*_1_, *a*_2_, *a*_3_ *and a*_4_:

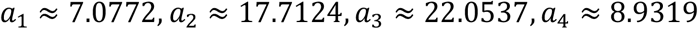

Therefore,

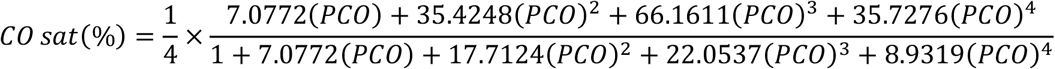

Since the data points are approximations and to make the final equation more practical for application, we round the resulting values, leading to the following equation.

After rounding the numbers, we get

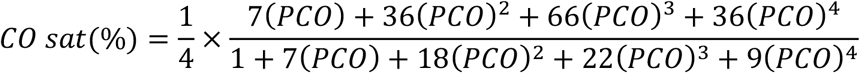

The unit of PCO is in ppt (parts per thousand)

The above two equations, when plotted, are almost indistinguishable.

From our original equation, we derive the following table, which demonstrates a good fit with the original research data [Hess et al., 2017].

**Table 1:**
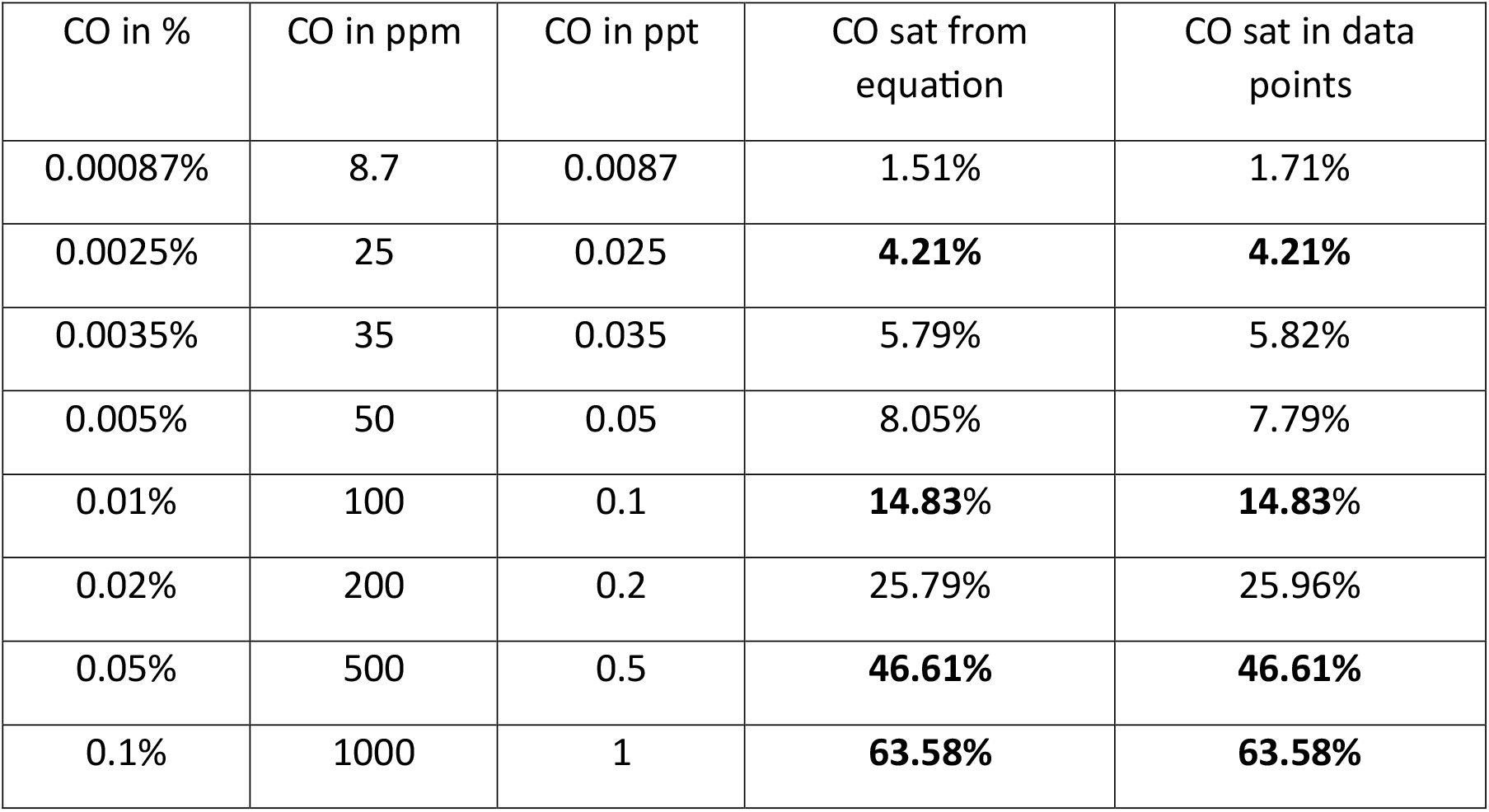
Carbon monoxide saturations at various partial pressures of carbon monoxide, demonstrating a strong fit between the equation’s results and published data.

The plot of our equation, shown in Fig. 1, exhibits a hyperbolic shape. This shape is consistent with and supports previous research findings [Douglas et al., 1912].

**Fig. 1:**
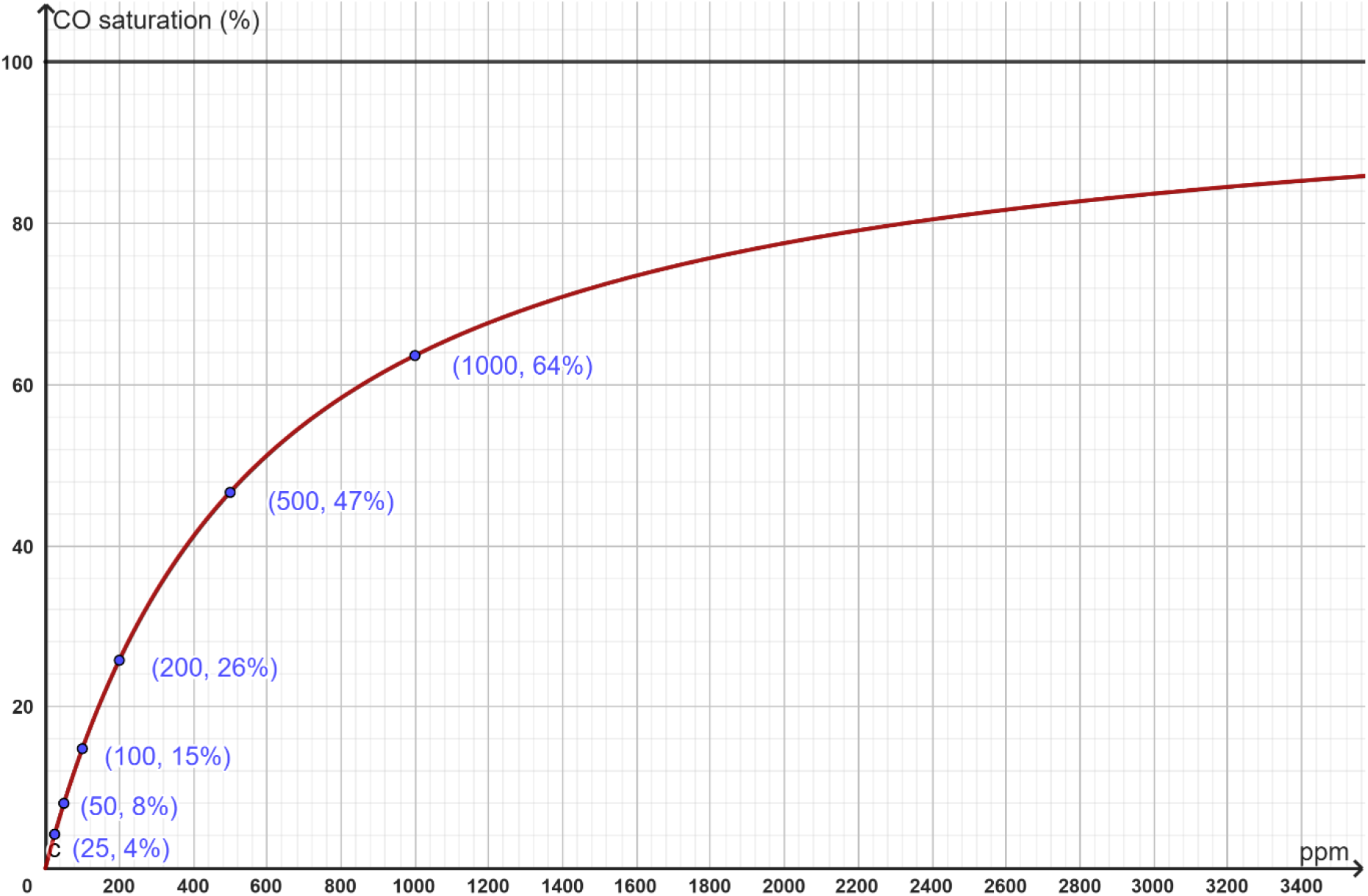
Plot of the CO saturation equation. The X-axis represents CO saturation (%) and the Y-axis represents CO pressure (ppm). Note the continuous decline in slope, a key characteristic of hyperbolic curves.

### Breakdown of each carboxyhemoglobin species

Unlike other equations in the literature, our equation allows us to determine the fractions of each of the five carboxyhemoglobin species. In each equation, the denominator represents the sum of all carboxyhemoglobin species, while the numerator corresponds to the specific carboxyhemoglobin species of interest.

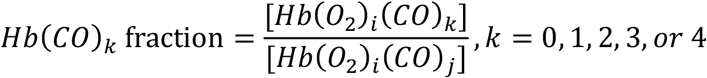

By comparing the above equation to the CO saturation equation,

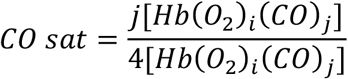

We can derive the following CO fraction equations.

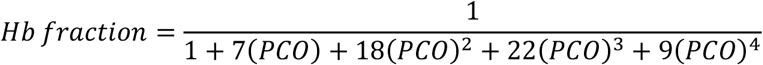

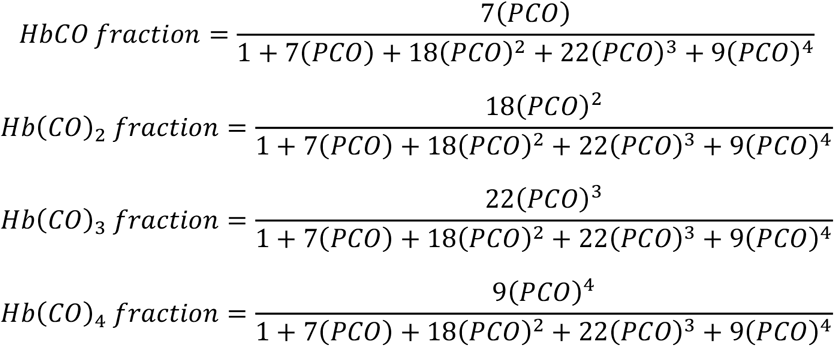

The plots of each fraction as a function of PCO, along with the superimposed CO saturation curve, are shown in Fig. 2.

**Fig. 2:**
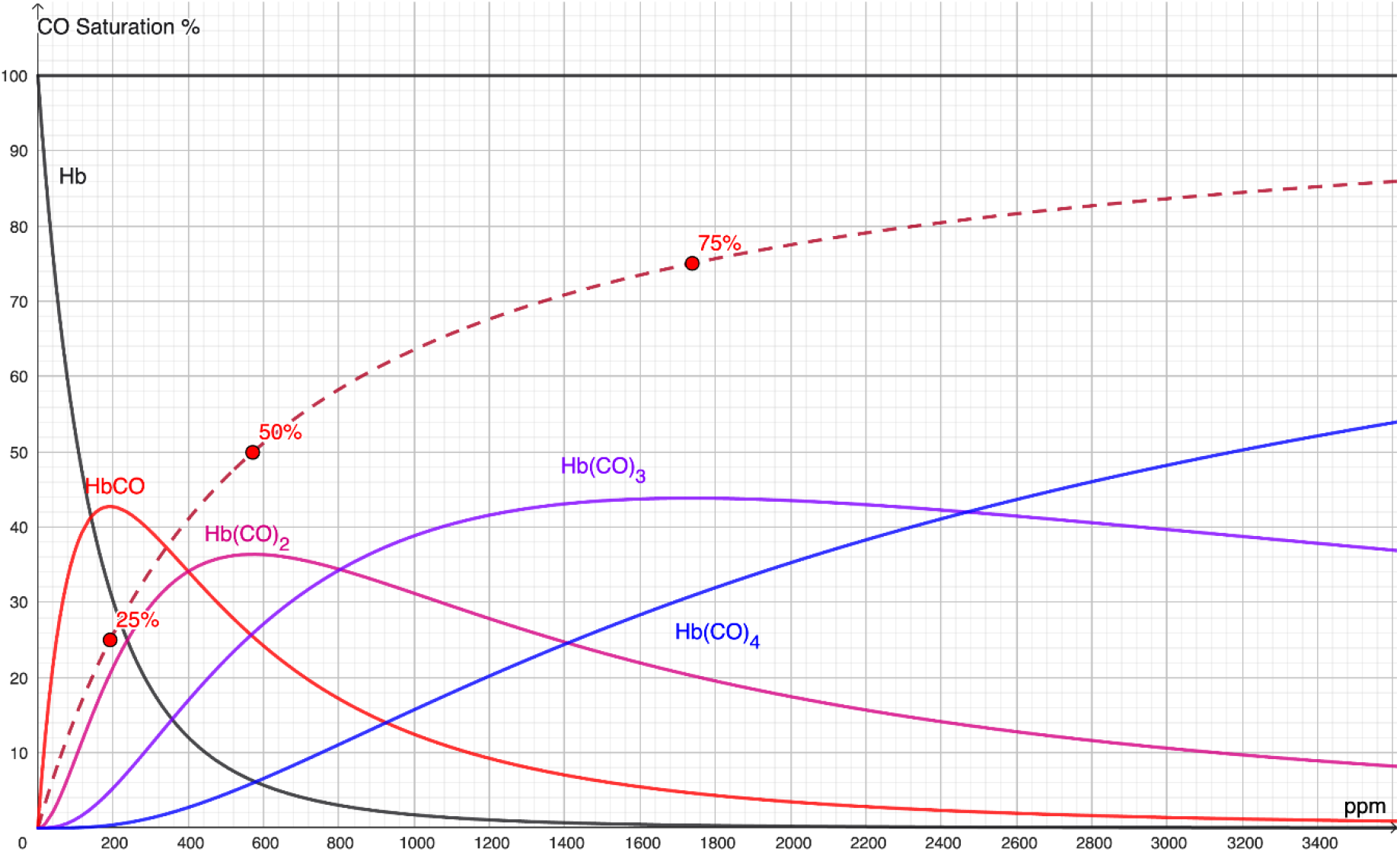
This graph displays the fraction equations for all carboxyhemoglobin species, with the carbon monoxide hemoglobin saturation curve superimposed. The graph shows that triply bound carboxyhemoglobin is the dominant species at pressures between 800 ppm and 2440 ppm. Notably, the fractions of Hb, HbCO, Hb(CO)_2_, Hb(CO)_3_, and Hb(CO)_4_ peak at the pressures corresponding to carboxyhemoglobin saturations of 0%, 25%, 50%, 75%, and 100%, respectively.

In Fig. 2, we observe that at minimal CO pressure, only a small amount of HbCO is present. As CO pressure increases, hemoglobin binds with carbon monoxide, forming singly bound carboxyhemoglobin. With further increases in CO pressure, additional carbon monoxide molecules displace oxygen molecules, leading to the formation of doubly bound carboxyhemoglobin. When CO pressure exceeds 800 ppm, triply bound carboxyhemoglobin becomes the dominant species. The fraction of quadruple-bound carboxyhemoglobin increases steadily, almost like a straight line, lacking the characteristic feature of cooperative binding.

It is interesting to note the contrast between oxygen binding and carbon monoxide binding to hemoglobin. In the case of oxygen binding, triply bound oxyhemoglobin has the lowest fraction, whereas for carbon monoxide binding, triply bound carboxyhemoglobin is much more abundant. Perrella et al. studied the intermediates of carbon monoxide and hemoglobin association [Perrella et al., 1992]. Because deoxyhemoglobin was used in their study, the distribution curves of the five carboxyhemoglobin species differ from those in our study [Perrella et al., 1992]. However, their curves allow us to visualize the fractions of singly, doubly, and triply bound carboxyhemoglobin species, which peak at CO saturations of 25%, 50%, and 75%, respectively. This feature is a unique characteristic of a rational saturation equation of the fourth degree, which can be proven mathematically. The laboratory work by Perrella et al. indirectly validates our equation.

## Discussion

The oxygen hemoglobin association equation was recently derived by Chou and Lee [Chou and Lee, 2023]. In the current paper, the author applies similar, albeit more complex, techniques to establish the carbon monoxide hemoglobin association equation.

Carbon monoxide is undetectable by humans; however, when present in higher concentrations, it can quickly cause respiratory distress. Buchelli et al. found that 20% of an unselected population has elevated carbon monoxide levels, defined as COHb ≥ 2.5% in nonsmokers and ≥ 5% in smokers [Buchelli et al., 2014]. Among those with elevated COHb, 78% are nonsmokers, with a mean level of 3.2%, while 22% are smokers, with a mean level of 6.7% [Buchelli et al., 2014]. Buchelli et al. determined that the primary source of CO exposure is likely from home environments.

Potential sources of carbon monoxide include volcanoes, bushfires, and automobile emissions. In clinical anesthesia practice, it is well known that desflurane and a desiccated CO_2_ absorbent (e.g., Baralyme) can produce carbon monoxide. Stabernack et al. reported CO concentrations as high as 36,000 ppm when desflurane, a commonly used anesthetic gas, flows through a desiccated CO_2_ absorbent like Baralyme [Stabernack et al., 2000]. Berry et al. reported a case of severe CO poisoning with an HbCO level of 36% during anesthesia with desflurane and Baralyme [Berry et al., 1999]. In that case report, no arterial CO pressure was provided. Using our equation, we can derive that the PCO would be 339.8 ppm.

The following table presents the concentrations of carbon monoxide under various conditions, along with the corresponding maximum saturations derived from our equation.

Notably, the pressure of CO in the Earth’s atmosphere is 100 ppb (parts per billion) [Hatheway et al., 2023]. For comparison, the atmospheric pressure of CO_2_ is significantly higher, currently at 412 ppm (0.0412%). Under normal conditions, our equation estimates the CO saturation to be 0.02%.

### Coburn-Forster-Kane (CFK) equation

The Coburn-Forster-Kane (CFK) equation, introduced in 1965, has been widely used to accurately estimate CO absorption, excretion, and saturation under various environmental CO concentrations, exposure durations, and alveolar ventilation rates [Coburn et al., 1965; Coburn, 2013; National Research Council, 2008]. However, the CFK equation does not directly describe CO saturation as a function of CO pressure. We believe our equation complements the CFK equation by providing additional insights into the behavior of carbon monoxide hemoglobin association, similar to the oxygen hemoglobin association.

The CFK equation can be expressed as the following differential equation: [Coburn et al., 1965, Coburn, 2013].

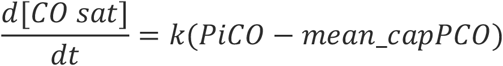

*PiCO*: inspired CO pressure; *mean*_*capPCO*: mean capillary CO pressure.

The value of *k* depends on endogenous CO production, blood volume, CO diffusion capacity, barometric pressure minus water vapor pressure, and alveolar ventilation [Coburn, 2013].

From the equation, we can see that the change in CO saturation approaches zero as the mean-capillary PCO nears the inspired PCO. Mathematically, this corresponds to the CO saturation curve reaching its maximum. Our data points are estimated from the CO saturation vs. CO exposure plot by Hess et al., where CO saturations have reached their maximum values [Hess et al., 2017].

The CFK equation describes CO saturation in relation to CO exposure. In contrast, our equation expresses CO saturation as a function of PCO at a PO_2_ level of 100 mmHg and, potentially, at any PO_2_ level. (See the section below: *CO Association Curves at Different Oxygen Pressures*.)

### The affinity of CO vs. O2 to hemoglobin

Historically, the following reaction has been used to investigate the relative affinity of hemoglobin for carbon monoxide versus oxygen.

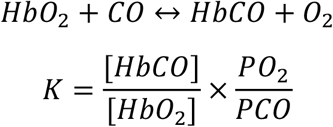

When the concentration of [HbCO] equals that of [HbCO_2_], the values cancel out, and past researchers have found that

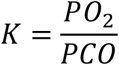

*K* was found to range from 200 to 490 [Benner, 2023; Fox, 1948; Delvau et al., 2020]. However, the chemical reaction and its corresponding equation shown above are inaccurate, as there are four distinct species of both oxyhemoglobins and carboxyhemoglobins. These notations [HbCO] and [HbO_2_] do not represent the whole story of the hemoglobin saturation of carbon monoxide or oxygen.

Using our equation, we can also calculate the ratio of 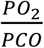 to determine the value of *K* at the point when both *CO* and *O*_2_ saturations are 50%. To do this, we first set the *CO* saturation equal to 50% and solve for *PCO*.

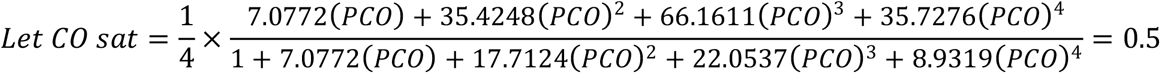

By solving the equation above, we get *PCO* ≅ 0.5724 *ppt*

Since CO saturation is set to 50%, it is reasonable to assume that O_2_ saturation is also 50% in this situation. Given that our equation is based on an equilibrium oxygen pressure of 100 mmHg, we obtain:

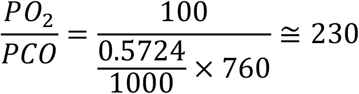

Thus, we derive and confirm the affinity of carbon monoxide for hemoglobin is 230 times greater than that of oxygen.

### Hyperbola shape vs. Sigmoidal shape

The carbon monoxide and hemoglobin association curve has been shown to be hyperbolic [Douglas et al., 1912], a finding that is confirmed by our equation. The hyperbolic shape in our graph reflects a continuous decline in slope, indicating that progressively larger increases in carbon monoxide concentration are required to achieve the same one percent increase in saturation.

In contrast, the sigmoidal shape oxygen and hemoglobin saturation curve concaves up first before concaving down, meaning that the hemoglobin can rapidly pick up oxygen molecules after the first *O*_2_ molecules initially bind at low oxygen pressure. The shape of the carbon monoxide association curve reflects the design of nature that our body rapidly uptakes the initial carbon monoxide molecules but resists the uptake of excessive carbon monoxide molecules.

### CO association curves at different oxygen pressures

We derive our equation at *PO*_2_ of 100 mmhg. Under different oxygen pressures, the equations will have different coefficients, resulting in different curves. As shown below, the coefficient *a*_*j*_ is a function of *PO*_2_.

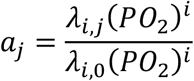

In theory, substituting different values of *PO*_2_ will yield a different set of *a*_*j*_ coefficient. and thus, different association curves. To compute these, we must first determine the values of all λs, which are considered constants—as suggested by the name, *association constants*.

### Solving for λs

Here, we propose a strategy to solve the general association equation.

1. At different levels of *PO*_2_, obtain four clinical data points of *PCO* and the corresponding carboxyhemoglobin saturations.
2. Use these data points to solve for the coefficients *a*_*j*_.
3. Once the *a*_*j*_ values are determined, solve for λ*s*. Here, we use *a*_1_ as an example.

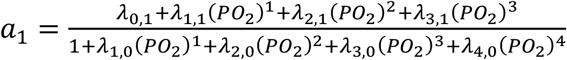
4. As shown in the equation above, we have eight λ values to solve for. Therefore, we need eight equations, each using different values of *a*_1_ and *PO*_2_
5. Repeat the same process for *a*_2_, *a*_3_ and *a*_4_

Once all the λ values are determined, we will derive the general equation of the carbon monoxide hemoglobin association curve. This general equation will have two variables: *PO*_2_ and *PCO*. By plugging in their values, we can calculate the *CO* saturation.

### Limitation

Although our approach is based on chemical kinetics and mathematical operations, it relies heavily on accurate data points. If different datasets are used, the resulting equation will differ. In this paper, we gathered data points from the literature. For future studies, arterial CO pressures and corresponding CO saturations can be measured in healthy individuals. These new data may yield an equation with different coefficients.

## Author Contributions

Chihjen Lee: conceptualization, methodology, data curation, data analysis, software, visualization, writing original draft, supervision.

Nikki Chen: data curation, validation, software, visualization, review and editing, proofreading

**Table 2:**
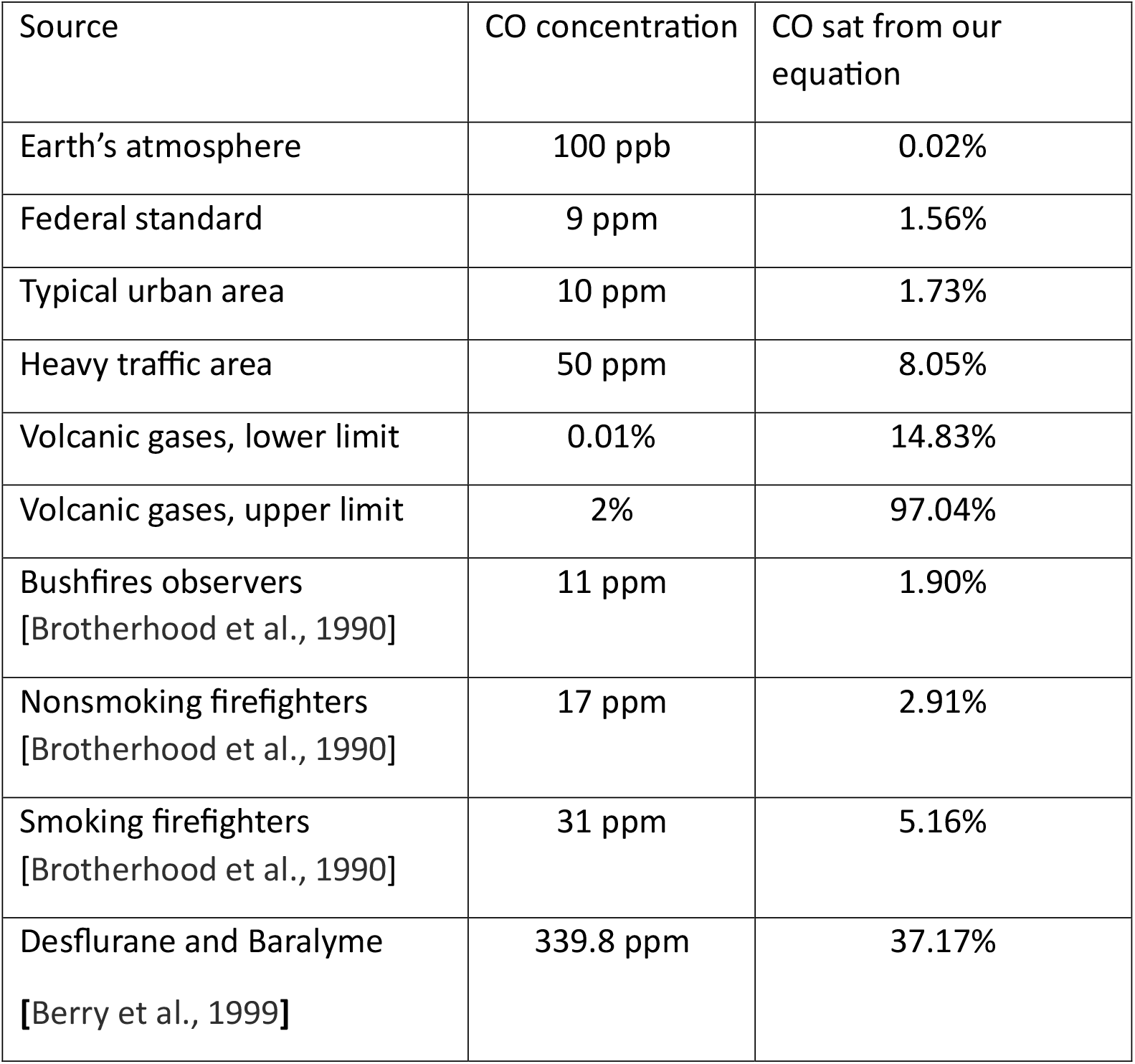
Carbon monoxide saturations calculated using our equation based on partial pressures in different environments.

## Notes

Conflict of interest: none

### Competing Interest Statement

The authors have declared no competing interest.

